# Simulated colonic fluid replicates the *in vivo* growth capabilities of *Citrobacter rodentium cpxRA* mutants and uncovers additive effects of Cpx regulated genes on fitness

**DOI:** 10.1101/2022.08.05.503015

**Authors:** Ashley Gilliland, Christina Gavino, Samantha Gruenheid, Tracy Raivio

**Author notes:** Address correspondence to Tracy Raivio.

## Abstract

*Citrobacter rodentium* is an attaching and effacing (A/E) pathogen used to model enteropathogenic and enterohemorrhagic *Escherichia coli* infections in mice. During colonization, *C. rodentium* must adapt to stresses in the gastrointestinal tract such as antimicrobial peptides, pH changes, and bile salts. The Cpx envelope stress response (ESR) is a two-component system used by some bacteria to remediate stress by modulating gene expression and is necessary for *C. rodentium* pathogenesis in mice. Here, we utilized simulated colonic fluid (SCF) to mimic the gastrointestinal environment which we show strongly induces the Cpx ESR and highlights a fitness defect specific to the Δ*cpxRA* mutant. While investigating genes in the Cpx regulon that may contribute to *C. rodentium* pathogenesis, we found that the absence of the Cpx ESR resulted in higher expression of the LEE master regulator, *ler*, and the genes *yebE, ygiB, bssR*, and *htpX* relied on CpxRA for proper expression. We then determined that CpxRA and select gene mutants were essential for proper growth in SCF when in the presence of extraneous stressors and in competition. Although none of the Cpx-regulated gene mutants exhibited marked virulence phenotypes *in vivo*, the Δ*cpxRA* mutant had reduced colonization and attenuated virulence, as previously determined, which replicated the *in vitro* growth phenotypes specific to SCF. Overall, these results indicate that the Δ*cpxRA* virulence defect is not due to any single Cpx regulon gene examined. Instead, attenuation may be the result of defective growth in the colonic environment resulting from the collective impact of multiple Cpx-regulated genes.

## Introduction

Attaching and effacing (A/E) pathogens like enteropathogenic and enterohemorrhagic *Escherichia coli* (EPEC and EHEC, respectively) are intestinal pathogens capable of causing severe diarrhea in children as well as adults (EHEC) (1). Due to the inability of EPEC and EHEC to replicate their human pathology in mice, *Citrobacter rodentium*, a murine A/E pathogen with genetic similarities to both EPEC and EHEC is used for *in vivo* models of infection (2, 3). *C. rodentium* causes colonic crypt hyperplasia in mice which results in severe inflammation in the colon (2, 4–6). Infections are lethal for some mice like strain C3H/HeJ, while C57Bl/6 mice experience self-limiting infections (7, 8). EPEC, EHEC, and *C. rodentium* all harbor the locus of enterocyte effacement (LEE) pathogenicity island that contains five operons (LEE1-5) encoding key proteins like intimin, Tir, and a type III secretion system (T3SS) which are responsible for the hallmark lesions formed upon intimate attachment to intestinal epithelial cells (1). The LEE is regulated by numerous factors including the master transcriptional activator Ler, encoded in the LEE1 operon, which activates the expression of LEE1-5. Ler also activates expression of the transcriptional activator, GrlA, and repressor, GrlR, encoded in the *grlRA* operon, which act upon LEE1 (5). In addition to LEE-encoded regulators, there are numerous other regulators of the LEE studied in EPEC and EHEC which activate or inhibit its expression based on environmental stressors sensed by the cell (9, 10).

A stress response that has been previously shown to be required for *C. rodentium* colonization and virulence is the Cpx envelope stress response (ESR) (11–13). The Cpx ESR is regulated by a two-component system (TCS) which consists of the sensor histidine kinase CpxA and the response regulator CpxR (14–16). In addition, it’s activity can be altered with two auxiliary signaling proteins, the inhibiting periplasmic protein CpxP and the activating outer membrane lipoprotein NlpE (16–18). While the sensing mechanism used by CpxA is poorly defined, inducing cues of the Cpx ESR include alkaline pH, salt, antimicrobial peptides, and misfolded inner membrane and periplasmic proteins (14, 16, 19–23). Upon activation, CpxA autophosphorylates and transfers a phosphoryl group to CpxR which then binds to a consensus sequence upstream of Cpx regulon members leading to altered expression. The CpxR binding consensus sequence 5’-GTAAA(N)_4-8_GTAAA-3’ is conserved to varying degrees however nucleotide deviations from the consensus sequence do not predict the relative expression of the downstream gene (20, 21, 24).

Previous studies investigating the role of the Cpx ESR in other pathogens have suggested mechanisms by which the Cpx ESR may impact, both negatively and positively, colonization and virulence. Some of these mechanisms include the negative regulation of *perC* resulting in reduced *ler* expression in EPEC, aiding the efficient expression of the bundle-forming pilus involved in initial host cell attachment in EPEC, and induction of genes encoding proteins required for maintaining envelope protein integrity and virulence regulation such as DegP, PpiA, and DsbA (9, 16, 25, 26). Overall, it has been concluded that the Cpx ESR facilitates adaptation to envelope stressors because it downregulates virulence genes and large protein complexes while upregulating envelope and protein modifying factors, although the specific reasons for its essentiality during pathogenesis have not been definitively demonstrated in most cases (16).

Previous work in *C. rodentium* has concluded that the Cpx ESR is activated *in vivo* based on the observation of increased expression levels of Cpx-regulated *cpxP* (11). Given that *C. rodentium* is an A/E pathogen which relies on the LEE and LEE-encoded T3SS for virulence, it is important to note that the absence of CpxRA does not impact the secretion of T3SS effector proteins *in vitro*, indicating its presence has a limited impact on overall LEE function (11, 13). In addition, inconsistent results have been reported on the ability of the Δ*cpxRA* mutant to grow in the virulence-inducing condition Dulbecco’s Modified Eagle Medium (DMEM) which is typically used as an *in vitro* growth condition prior to conducting *in vivo* models. Thomassin et al. (11) measured an extended lag phase for cells lacking CpxRA, while Vogt et al. (13) found no growth defect in the same media. To further investigate the role of the Cpx ESR in pathogenesis, the Cpx regulon of *C. rodentium* was defined by microarray, RNA-seq, and SILAC proteomic analysis (12, 13). The data produced from these two studies was extensive and the impacts of numerous genes of interest on virulence have not been investigated. Giannakopoulou et al. (12) determined that the auxiliary proteins, CpxP and NlpE, were not required for colonization or virulence in *C. rodentium*. On the other hand, Vogt et al. (13) showed that the Cpx regulon members *degP* and *dsbA* were required for *C. rodentium* virulence in C3H/HeJ mice, however the Cpx regulation of these genes was not responsible for the virulence defect seen when *cpxRA* was absent. The Cpx regulon is extensive with the presence of CpxRA contributing to the differential expression of over 330 transcripts in *C. rodentium* (12, 13). The roles of some of the more strongly upregulated genes in both studies, such as *yebE, ygiB, bssR* and *htpX* which are investigated in this study, are not defined in *C. rodentium*.

In this study, we aimed to elucidate the role of the Cpx ESR in *C. rodentium* colonization and virulence utilizing simulated colonic fluid (SCF), first developed by Beumer et al. (27). Here we show that SCF is a preferable medium for *C. rodentium* growth, highlights previously unobserved growth defects in Cpx-regulated gene mutants, and more accurately predicts a mutants’ performance *in vivo* relative to LB and HG-DMEM. Using luminescent reporters, generating single, double, and triple mutants, and testing strains in competition with fluorescent wild-type *C. rodentium*, our data suggests the Cpx ESR is required for adequate survival and proliferation in the gut environment, modelled by SCF, by ameliorating stressors and ensuring the appropriate regulation of virulence and envelope-associated genes which contributes to the colonization and pathogenesis of *C. rodentium*.

## Materials and Methods

### Bacterial Strains and Growth Conditions

Bacterial strains used are listed in Table S1. Unless otherwise indicated, cells were grown in lysogeny broth (LB; 10 g/L tryptone, 5 g/L yeast extract, and 5 g/L NaCl), with appropriate antibiotics at 37°C and 225 rpm. Cells were also grown on LB with 1.5% agar plates and incubated at 37°C for 16-18 hours. When required, media was supplemented with: 30 ug/ml kanamycin, 100 ug/ml ampicillin, 25 ug/ml chloramphenicol, 0.3mM diaminopimelic acid, 5% sucrose, 0.2 ug/ml anhydrotetracycline.

### Deletion Mutant Construction

All deletion mutants were generated using allelic exchange in the methods described by Vogt et al. (13). In summary, in-frame deletion constructs were generated using overlap extension PCR and the primers listed in Table S2. Amplicons were digested using the XbaI and SphI/PaeI, ligated into pUC18, and transformed into OneShot TOP10 chemically competent cells (Invitrogen, USA). Plasmids were mini-prepped and sent for Sanger sequencing for amplicon confirmation. Once confirmed, the deletion construct was digested out of pUC18 and ligated into the suicide vector, pRE112, where it was transformed by electroporation into MFDpir cells (28). MFDpir cells containing the deletion construct underwent conjugation with a recipient strain of *C. rodentium* DBS100, and single-crossover colonies were selected with chloramphenicol. Single-crossover colonies were confirmed by PCR using primers designed to flank the deletion site by ∼75 bp on each side (Table S2). Loss of pRE112 was determined by plating on LB -NaCl with 5% sucrose (filter-sterilized) and growing at room temperature for 2 days. Colonies that were sucrose-resistant and chloramphenicol-sensitive were screened by PCR to confirm intended deletion.

### Luminescent Reporter Construction

Luminescent reporters were constructed using the pNLP10 *lux-*reporter plasmid (21). Primers listed in Table S2 were designed to amplify ∼500bp upstream and 50 bp downstream of the translation start site apart from *ygiB*, where the amplified promoter was ∼300 bp. The restriction enzymes EcoRI and BamHI were incorporated into the forward and reverse primers respectively, unless otherwise indicated. Promoter regions were amplified, cloned into pNLP10 and transformed into OneShot TOP10 chemically competent cells (Invitrogen, USA). TOP10 colonies harboring the pNLP10 plasmid were confirmed for insert presence using colony PCR with primers flanking the pNLP10 multiple cloning site as well as with Sanger sequencing. Plasmids with the correct insert were mini-prepped and transformed into electrocompetent *C. rodentium* wild-type or mutant cells.

### Luminescence Assays

Strains grown overnight in biological triplicate were subcultured 1:100 and grown to an optical density (OD_600_) of 0.4 – 0.6. Following this, subcultures were divided into 1 ml aliquots, centrifuged and the subculturing media was removed. Cells were then resuspended in 1 ml of fresh media containing 30 ug/ml kanamycin (T = 0). Resuspension media included: LB, high-glucose Dulbecco’s Modified Eagle Medium with no phenol (HG-DMEM) (Gibco™, cat. no. 31053028), and simulated colonic fluid, using ox bile in lieu of porcine bile (SCF: 6.25 g/L proteose-peptone, 2.6 g/L glucose, 0.88 g/L NaCl, 0.43 g/L KH_2_PO_4_, 1.7 g/L NaHCO_3_, 2.7 g/L KHCO_3_, 4.0 g/L ox bile, and 0.1M MOPS) (27). HG-DMEM and SCF were buffered with 0.1M MOPS to maintain a physiological pH of 7.4 and 7.0 respectively, unless otherwise indicated. SCF was prepared fresh for each experiment and filter sterilized. For luminescent assays with sub-inhibitory levels of oxidative stress, H_2_O_2_ was added to a final concentration of 1mM in LB and SCF, both buffered with 0.1M MOPS.

200 ul of induced cells were then transferred into a black-walled clear bottom 96-well plate and incubated at 37°C shaking, unless otherwise indicated. Empty wells were left between strains to prevent contaminating luminescence from adjacent wells. OD_600_ and luminescence measurements were taken either 1-hour post-resuspension or over 6 hours starting at 0.5-hour post-resuspension using the Victor X3 2030 multilabel plate reader (Perkin Elmer). Luminescence was measured in counts per second (CPS) and normalized using the measured OD_600_ of the same well to accommodate for differences in cell number between cultures. Prior to calculating CPS/OD_600_, the OD and CPS values measured for blank wells containing each induction medium were subtracted from experimental wells. Biological triplicates were averaged to determine the mean luminescence in each condition and statistical significance was calculated using a Student’s *t-*Test. For *cpxP-lux* activity in mutant strains, luminescence was compared to wild-type levels of *cpxP-lux* using a one-way ANOVA with Dunnett’s multiple comparison test. Luminescence assays were repeated at least twice with the data from one representative experiment shown.

### Growth Curves

Strains were grown in biological triplicate overnight, washed with 1X PBS, and standardized to an OD_600_ of 1.0. Cells were inoculated 1:100 into media aliquoted in a clear 96-well plate. Growth media included LB, HG-DMEM, SCF, and SCF with no MOPS. Growth assays conducted in the presence of oxidative stress were prepared in the same manner with the addition of hydrogen peroxide to a final concentration of 1mM. For these assays, both the LB and SCF was buffered with 0.1M MOPS to control for pH changes in both medium. Plates were read in an Epoch2 microplate reader (Biotek, USA) set to 37°C with continuous orbital shaking at a frequency of 237 cpm (4mm) at slow speed. Blank wells were subtracted from corresponding culture wells prior to calculations. Biological triplicates were averaged with data points indicating the mean and the standard deviation indicated by error bars. Growth curves were completed at least twice with the data from one experiment shown.

### *Construction of Fluorescent* amCyan C. rodentium

A *C. rodentium* strain containing a chromosomal, tetracycline-inducible, a*mCyan* was constructed via two sequential rounds of allelic exchange. The purpose of the first round of allelic exchange was to integrate *tetRA* genes into the *C. rodentium* gene *xylE. xylE* has been used in previous studies as an integration site for constitutive *lux* expression as well as tetracycline-induced expression of *tccP*, and its interruption has been shown to not impair *C. rodentium* fitness (29, 30). Using the primers listed in Table S2, fragments 1 and 3 constituting the intended flanking regions of *xylE* were amplified from *C. rodentium* DBS100 genomic DNA based off the primers designed by Girard *et al*. (30), while fragment 2 containing the *tetRA* genes was amplified from the *E. coli* commensal MP7 (31), which contains a chromosomal tetracycline*-*inducible *mcherry*. The three fragments were joined 1-2-3 using overlap-extension PCR and amplified with primers containing restriction sites for SphI and SacI creating the fragment *xylE-tetRA-xylE*. Confirmation of the fragment, generation of the donor strain and conjugation was then carried out as previously described for construction of deletion mutants. Sucrose-resistant, tetracycline-resistant, and chloramphenicol-sensitive *C. rodentium* colonies were PCR screened to confirm insertion of *tetRA* into *xylE* (DBS100-*tetRA*).

To insert the *amCyan* gene downstream of *tetRA* and using the primers listed in Table S2, fragment 1 containing the 3’-end of *tetA* was amplified from MP7 (31), fragment 2 containing *amCyan* was amplified from the plasmid pFU78 (32), and fragment 3 containing the portion of *xylE* homologous to that which is downstream of the *tetA* gene in the strain DBS100-*tetRA* was amplified from DBS100 genomic DNA. The three fragments were fused 1-2-3 using overlap extension PCR to generate the new fragment *tetA-amCyan-xylE* which was then sequenced in pUC18, sub-cloned into pRE112, and transformed into MFDpir for conjugation with DBS100-*tetRA*. Successful transconjugants were selected for on LB -NaCl with 5% sucrose plates containing tetracycline. After 2 days of growth at room temperature, plates were imaged using the ChemiDoc system, using Cy2 for visualization of fluorescent *amCyan C. rodentium* colonies (DBS100-AC) which were then selected for further confirmation via PCR.

### Competitive Assays

*In vitro* competitive assays between DBS100-AC and the generated mutant strains were conducted in both LB and SCF following the protocol previously described in Pick *et al*. (33), with minor changes. In summary, overnight cultures of each strain grown in biological triplicate were standardized to an OD_600_ of 1.0. The Δ*cpxRA* mutant was standardized to an OD_600_ of 1.4 as previous enumeration indicated overnight cultures had less viable cells relative to the other strains (data not shown). 10 ul of DBS100-AC and 10 ul of the competitor strain were then inoculated into 2 ml of either LB or SCF. Cocultures were vortexed and a 100 ul aliquot was removed for serial dilution which was plated using the track dilution method on LB agar containing 0.2 ug/ml anhydrotetracycline (34). The cocultures were immediately incubated at 37°C for 24 hours following aliquot removal, while plates with track dilutions were incubated at 30°C overnight. Plates were imaged using the ChemiDoc system, using Cy2 for visualization of fluorescent DBS100-AC. The number of competitor (non-fluorescent) colonies was determined by subtracting the number of fluorescent colonies from the total colony count. At 24 hours, cocultures were subcultured 1:100 into fresh media and an aliquot from the mature coculture was serial diluted and plated for enumeration. At 48 hours, the mature coculture was serial diluted and plated for enumeration.

### C57Bl/6J and C3H/HeJ Mouse Infections

All animal experiments were performed under conditions specified by the Canadian Council on Animal Care and were approved by the McGill University Animal Care Committee. C57BL/6J and C3H/HeJ mice were purchased from the Jackson Laboratory (Bar Harbor, ME, USA). Five-week-old female mice were infected by oral gavage with 0.1 ml of LB medium containing 2-3 × 10^8^ colony-forming units (CFU) of bacteria. The infectious dose was verified by plating serial dilutions of the inoculum onto MacConkey agar (Difco). For survival analysis of C3H/HeJ mice, the mice were monitored daily and were killed if they met any of the following clinical endpoints: 20% body weight loss, hunching and shaking, inactivity, or body condition score of <2 (35). To monitor bacterial colonization, fecal pellets or the terminal centimeter of the colon were homogenized in PBS, serially diluted, and plated on MacConkey agar. Plates containing between 30 and 300 colonies were counted. Spleens were removed and weighed, and splenic indexes were calculated [v(weight of spleen × 100/weight of mouse)].

## Results

### Simulated colonic fluid induces the Cpx envelope stress response

Previous work in the field has utilized HG-DMEM to induce virulence gene expression and test mutant fitness *in vitro* prior to conducting *in vivo* experiments. To further investigate the colonization defect associated with the Δ*cpxRA* mutant *in vitro*, we used the medium SCF, meant to mimic gastrointestinal conditions, which includes relevant components like bile.

Initially, it was important to determine whether the Cpx ESR was active in SCF. A luminescent reporter for the Cpx regulon member *cpxP* was used to measure Cpx ESR activation in both wild-type and Δ*cpxRA* cells grown in LB, HG-DMEM, and SCF. Our results indicate that both HG-DMEM and SCF strongly induce the Cpx ESR in wild-type cells (Figure 1A-B). Meanwhile in the Δ*cpxRA* mutant, *cpxP-lux* was not detectable in any of the conditions used in this study (Figure 1A, Figure S1). Unexpectedly, the *cpxP-lux* expression pattern over time and by extension, the activity of the Cpx ESR, differed depending on the medium used for growth thus indicating condition-specific activity levels (Figure 1B). In addition, while having a less substantial impact on the activity of the Cpx ESR, cultures that were grown in LB shaking had higher levels of *cpxP-lux* activity relative to static cultures, whereas cultures grown statically in HG-DMEM and SCF had higher *cpxP-lux* activity compared to shaking cultures (Figure 1C-E). These results suggest that numerous factors including media, growth phase, and culture conditions all contribute to the activity of the Cpx ESR and should be considered when investigating the activity of envelope stress responses *in vitro*.

**Figure 1.**
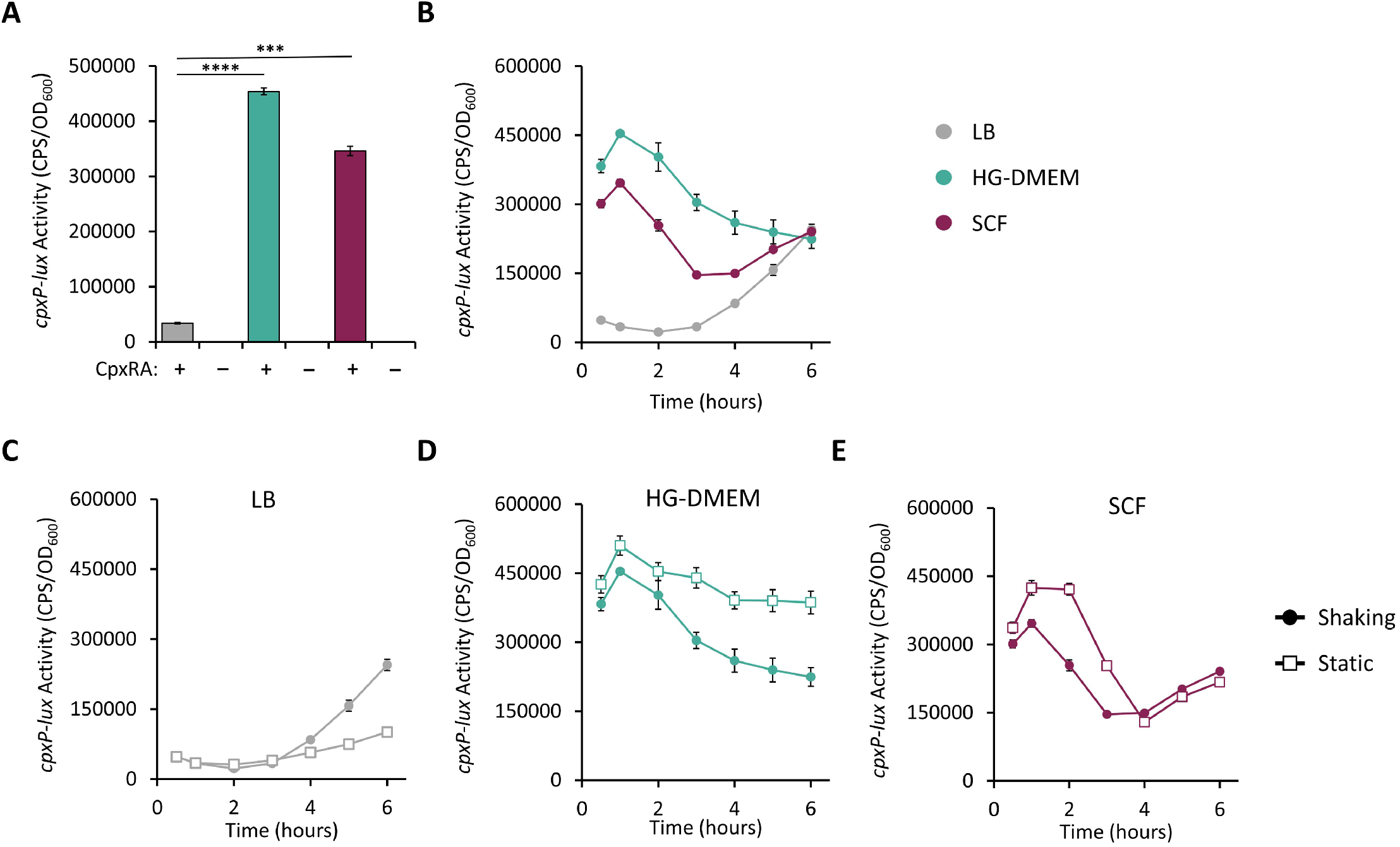
The Cpx ESR is induced in HG-DMEM and SCF. Luminescent assays indicating the activity of *C. rodentium cpxP-lux* in LB (gray), HG-DMEM (teal), and SCF (maroon). (A) Activity of *cpxP-lux* promoter in wild-type (+) and Δ*cpxRA* (–) cells measured 1-hour post-resuspension (\*\*\**P* < 0.001, \*\*\*\**P* < 0.0001, Student’s *t*-test). (B) Luminescence generated from wild-type *cpxP-lux* promoter over time starting from 0.5-hour post-resuspension. (C-E) Activity of wild-type *cpxP-lux* promoter measured over time from cells grown in either shaking (filled circle) or static (open square) conditions starting from 0.5-hour post-resuspension. Luminescence was measured in counts per second (CPS) and normalized to bacterial optical density (OD_600_). Data points represent the mean of three biological replicate cultures with error bars indicating standard deviation.

### *LEE master regulator*, ler, *expression responds to SCF and HG-DMEM*

Previous studies have demonstrated the *C. rodentium* Δ*cpxRA* mutant does not have a significantly altered T3SS secreted protein profile *in vitro* suggesting the LEE-encoded T3SS remains functional (11, 13). On the other hand, in EPEC and EHEC there is reduced transcription of LEE operons upon activation of CpxRA as well as reduced expression and secretion of translocator proteins (36, 37). Using a *ler-lux* reporter, we investigated whether the attenuation of the *C. rodentium* Δ*cpxRA* mutant could be in part due to mistimed or inappropriate expression levels of the LEE master regulator *ler*. Expressions levels were measured in SCF, to mimic the colonic lumen, and tissue-culture medium HG-DMEM, as a proxy for the intestinal epithelial cell layer. Interestingly, wild-type and Δ*cpxRA* mutant cells grown in SCF exhibited similar low expression levels of *ler-lux* (Figure 2A). Meanwhile, the Δ*cpxRA* mutant had significantly increased *ler-lux* expression when grown in HG-DMEM (Figure 2B). In addition, our results indicate that *ler-lux* expression is more strongly activated in HG-DMEM relative to SCF which supports the notion that LEE gene expression is dependent on the surrounding environment and likely has increased expression at the epithelial cell surface relative to the colonic lumen (Figure 2). Experiments were repeated in static conditions as well as using low-glucose DMEM (LG-DMEM) where the increased levels of *ler-lux* expression could also be observed in the Δ*cpxRA* mutant (Figure S2). In culmination, these results support the hypothesis that the Cpx ESR is an important environmental sensor which contributes to the timing and extent of expression of the LEE, which could contribute to the attenuation of the Δ*cpxRA* mutant.

**Figure 2.**
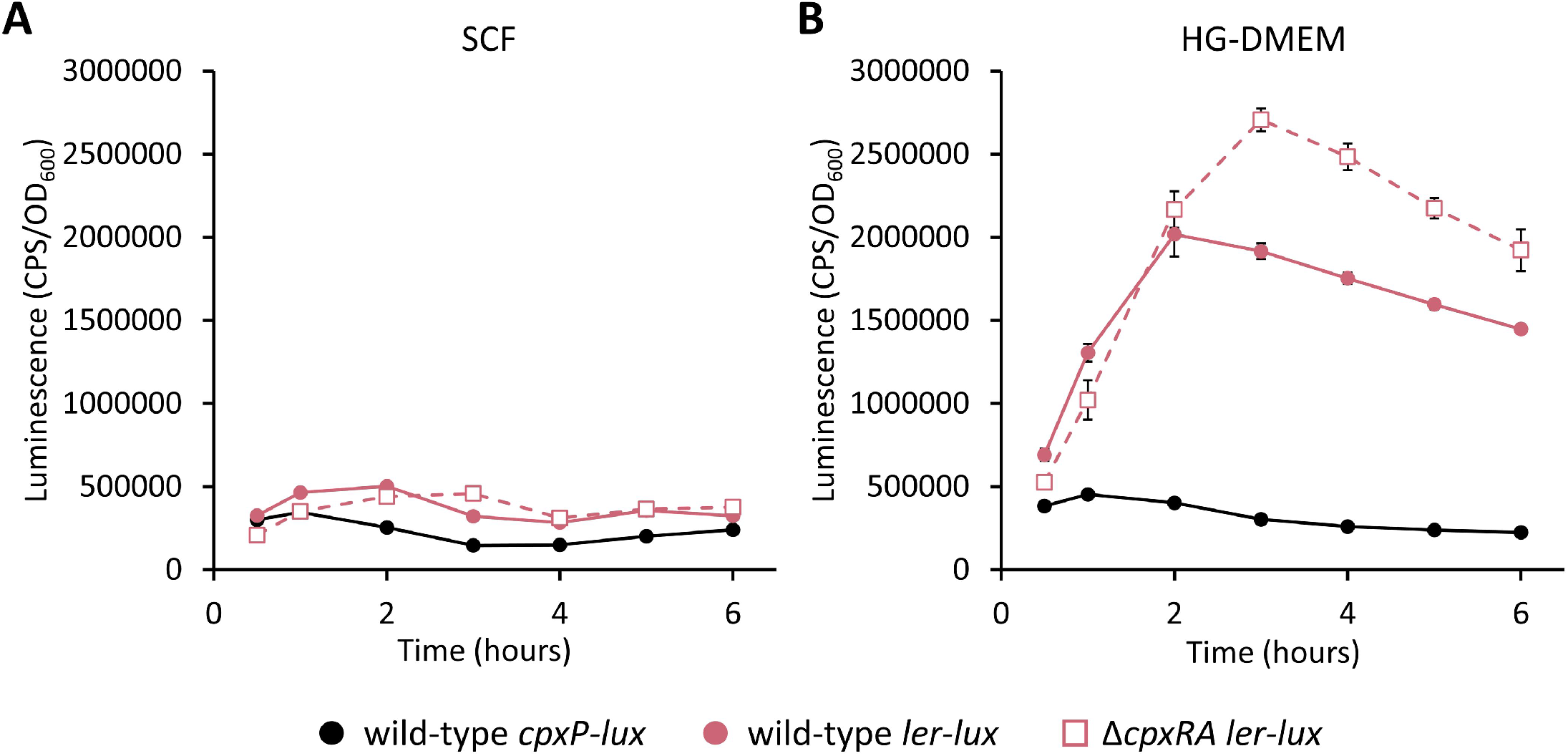
The impact of the Cpx ESR on the expression of the major LEE regulator, *ler*, is greater in HG-DMEM relative to SCF. Luminescent assays indicating the activity of *C. rodentium cpxP-* (black) and *ler-lux* (pink) reporters in wild-type (closed circle) and Δ*cpxRA* (open square) cells grown in (A) SCF or (B) HG-DMEM. Activity was measured over time starting from 0.5-hour post-resuspension. Luminescence was measured in counts per second (CPS) and normalized to bacterial optical density (OD_600_). Data points represent the mean of three biological replicate cultures with error bars indicating standard deviation.

### Identification and confirmation of Cpx regulon members with potential virulence contribution

Research done by two independent groups collected proteomic, RNA-Seq and microarray data to identify genes that were differentially expressed in the absence of CpxRA (12, 13). Using the data collected, the genes listed in Table 1 were selected for further study based on their predicted function, expression levels, and/or lack of previous investigation. *htpX* and *yebE* had some of the highest changes in transcript abundance when the *cpxRA* locus was ablated besides *cpxP* and *yccA*, which have both been previously investigated in *C. rodentium* (13). The SILAC data for *htpX* was insignificant due to the detection of only 1 peptide while *yebE* had a 5.36-fold higher peptide count in wild-type cells with a *P-*value of 0.08. *ygiB* had significantly higher transcript abundances in both transcriptomic studies as well as peptide counts in the presence of CpxRA as indicated by bolded values (Table 1). *bssR* is involved in biofilm regulation and had significantly higher transcript abundance in wild-type cells in both transcriptomic studies despite an absence of detection in the SILAC data (Table 1).

**Table 1.** Mined RNA-Seq and SILAC data from Vogt *et al*. (13) and microarray data collected in Giannakopoulou *et al*. (12).

| Data Source | Dataset S3- Vogt *et al.* (2019) |  | Giannakopoulou *et al.* (2018) | Protein Function<sup>3</sup> |
| --- | --- | --- | --- | --- |
| Gene Name | Fold change RNA-Seq<sup>1</sup> | Fold change SILAC<sup>1</sup> | log2 fold change $\Delta cpxRA$-WT microarray<sup>2</sup> | |
| *htpX* | **12.10** | 10.77 | **-0.93** | Membrane-localized protease |
| *yebE* | **24.52** | 5.36 | -1.92 | Inner membrane protein with transmembrane domain |
| *ygiB* | **3.54** | **2.24** | **-0.99** | Outer membrane protein |
| *bssR* | **3.72** | n.d. | **-3.36** | Biofilm regulator involved in catabolite repression and stress response |

**Bolded** values indicate $P_{adj} < 0.05$ (RNA-Seq), $FDR < 0.05$ (SILAC), $t$ -test $< 0.05$ (microarray).
<sup>1</sup>Calculated [wild-type/ $\Delta cpxRA$ ]
<sup>2</sup>Negative value indicates higher expression in wild-type DBS100
<sup>3</sup>Protein function as described by UniProtKB for *E. coli* K12
<sup>4</sup>Included due to potential operon leader of gene of interest

Using the *lux-*reporter plasmid pNLP10, four reporters were constructed and tested in LB broth, the virulence-inducing media HG-DMEM, and SCF (21, 38, 39). While some of these genes, namely *yebE* and *htpX*, have had Cpx-dependent expression confirmed experimentally in previous studies using other bacterial strains, their expression has never been studied in *C. rodentium* DBS100 (20, 21, 40, 41). Here we show that the expression of *yebE, ygiB, bssR*, and *htpX* relies on the Cpx ESR in LB, HG-DMEM, and SCF (Figure 3A-D). Furthermore, all four reporters had significantly increased expression in HG-DMEM and SCF relative to the expression levels measured in LB in a Cpx-dependent manner (Figure 3A-D). When comparing the expression of the four reporters in the Δ*cpxRA* mutant background, only *htpX-lux* indicated differences between the media tested (Figure S3). The *yebE-lux* reporter indicated the greatest level of induction as promoter activity was upregulated 59- and 29-fold in HG-DMEM and SCF, respectively, relative to the measured expression in LB (Figure 3A). Finally, each reporter gene, except for *yebE*, had increased luminescence in wild-type cells when grown on solid LB agar relative to Δ*cpxRA* mutant cells (Figure S4A).

**Figure 3.**
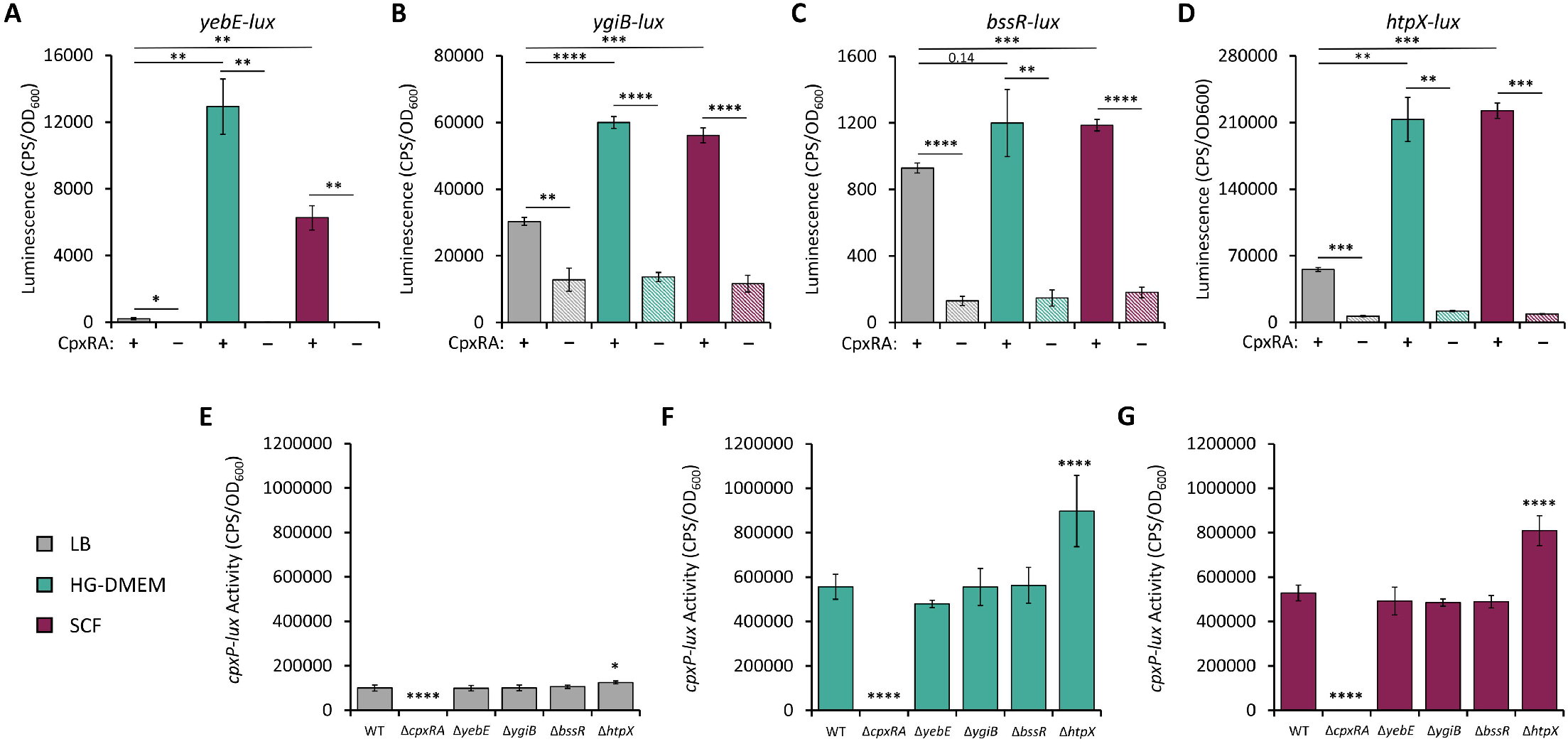
Cpx-regulated genes have increased expression in HG-DMEM and SCF while the activity of the Cpx ESR is only impacted in Δ*htpX* cells. (A-D) Luminescent assays indicating the promoter activity of *C. rodentium* genes of interest in LB (gray), HG-DMEM (teal), and SCF (maroon). Activity of (A) *yebE-*, (B) *ygiB-*, (C) *bssR-*, and (D) *htpX-lux* promoters in wild-type (+, solid bars) and Δ*cpxRA* (–, striped bars) cells measured 1-hour post-resuspension. Luminescence was measured in counts per second (CPS) and normalized to bacterial optical density (OD_600_). Bars represent the mean of three biological replicate cultures with error bars indicating standard deviation (\**P* < 0.05, \*\**P* < 0.01, \*\*\**P* < 0.001, \*\*\*\**P* < 0.0001, Student’s *t*-test). (E-G) Mutant strains harboring *cpxP-lux* reporter plasmids were grown in in (E) LB, (F) HG-DMEM, and (G) SCF. Bars represent the mean of three biological replicate cultures with error bars indicating standard deviation. The asterisks indicate a statistically significant difference between the mutant strain relative to wild-type cells (\**P* < 0.05, \*\*\*\**P* < 0.0001, one-way ANOVA with Dunnett’s multiple comparison test).

Given that our evidence indicates *yebE, ygiB, bssR*, and *htpX* rely on the presence of CpxRA for proper expression, we questioned whether the absence of these genes would impact the activity of the Cpx ESR. Using allelic exchange, *C. rodentium* knockout mutants were generated for *yebE, ygiB, bssR*, and *htpX* and transformed with the *cpxP-lux* reporter to measure Cpx activity in LB, HG-DMEM, and SCF. Our results indicate that only Δ*htpX* cells had significantly increased expression of *cpxP* in LB, HG-DMEM, and SCF, supporting previously reported findings observed in *E. coli* K-12 strain MC4100 (Figure 3E-G) (40). Increased *cpxP-lux* expression was also seen on LB agar plates (Figure S4B).

### *Cpx-regulated genes impact* C. rodentium *fitness in simulated colonic fluid*

Previous research has shown differing results with regard to the effect of removing the Cpx ESR on *C. rodentium* growth *in vitro*. Vogt et al. (13) showed no growth defects associated with a *ΔcpxRA* mutant in shaking LB broth or static high-glucose DMEM with 5% CO_2_. On the other hand, Thomassin et al. (11) found that *C. rodentium* Δ*cpxRA* cultures had a longer lag phase in DMEM but eventually would grow to a comparable OD to the wild-type and complemented strains. In this study, all the mutants tested grew comparably in LB (Figure 4A). In HG-DMEM, the mutants grew slower and to a lower OD relative to wild-type cells (Figure 4B). One predominant issue to note with *C. rodentium* growth in HG-DMEM is that wild-type cells can neither grow quickly or to high density as seen by the exponential growth phase starting at approximately 3.5 hours and ending at 10.5 hours while only reaching an OD_600_ maximum of 0.2 (Figure 4B). Contrary to this, when grown in SCF, *C. rodentium* wild-type cells grew rapidly and reached an OD of 0.7 at 6.5 hours (Figure 4C). Interestingly, only the Δ*cpxRA* mutant had a marked growth defect when grown in SCF relative to the other strains (Figure 4C). Furthermore, when SCF was not buffered and therefore unable to maintain a pH of 7 over time, the Δ*cpxRA* mutant could not grow, suggesting that unstable pH is toxic to these cells (Figure 4D). A *cpxRA* complemented strain had restored growth and Cpx activity, measured by expression of a *cpxP-lux* reporter, in all conditions tested (Figure S5). While not exhibiting growth defects in buffered SCF (Figure 4C), in unbuffered SCF both the Δ*ygiB* and Δ*htpX* mutants experienced an extended lag phase as well as had increased variability between biological replicates highlighting growth defects associated with these mutants (Figure 4D).

**Figure 4.**
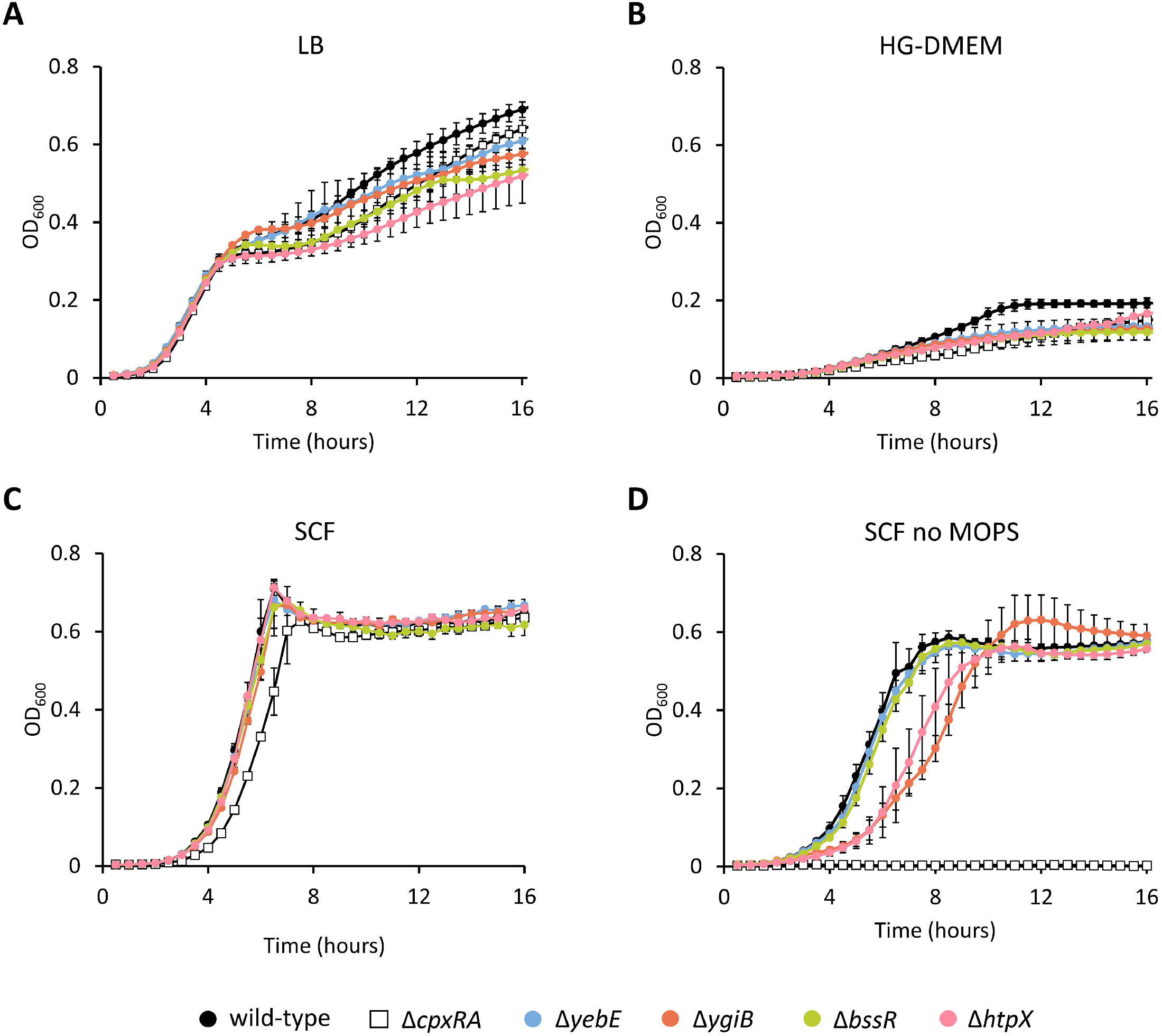
SCF promotes *C. rodentium* growth and highlights fitness defects in Δ*cpxRA*, Δ*ygiB*, and Δ*htpX* cells. Overnight cultures were standardized to an OD_600_ of 1.0 then subcultured 1:100 into (A) LB, (B) HG-DMEM, (C) SCF, and (D) SCF without MOPS buffer. Data points represent the mean of three biological replicate cultures with error bars indicating standard deviation.

Beyond unstable pH, another stressor relevant to the gastrointestinal tract and *C. rodentium* proliferation is the presence of oxidative stress. To test this physiologically relevant stressor, we grew the wild-type and mutant strains in MOPS-buffered LB and SCF with a sub-inhibitory level of hydrogen peroxide. In LB, the Δ*cpxRA* mutant had an extended lag phase relative to the other strains, although it reached a comparable OD after 9 hours growth (Figure 5A). Interestingly, similar to the growth in unbuffered SCF shown in Figure 4D, the Δ*cpxRA* mutant was unable to grow in buffered SCF containing 1mM hydrogen peroxide (Figure 5B). Growth was restored to wild-type levels in the *cpxRA* complemented strain (Figure S5E). In addition, the mutants Δ*yebE*, Δ*ygiB*, and Δ*htpX* all exhibited varying degrees of susceptibility to sub-inhibitory levels of hydrogen peroxide in SCF, with Δ*ygiB* and Δ*htpX* mutants having the most severe growth defects (Figure 5C-F). Finally, a consistent reduction in OD indicating cell lysis was also observed in SCF as cells transitioned from exponential to stationary phase (Figure 4C, Figure S5C and E, Figure 5B-F). The cause of this reduction requires further study. These results demonstrate that SCF is a suitable growth medium for *C. rodentium* as indicated by the observed rapid exponential phase, high OD_600_, and stable stationary phase. In addition, the Δ*cpxRA* mutant is extremely sensitive to physiologically relevant stressors present in the gastrointestinal environment like unstable pH and oxidative stress.

**Figure 5.**
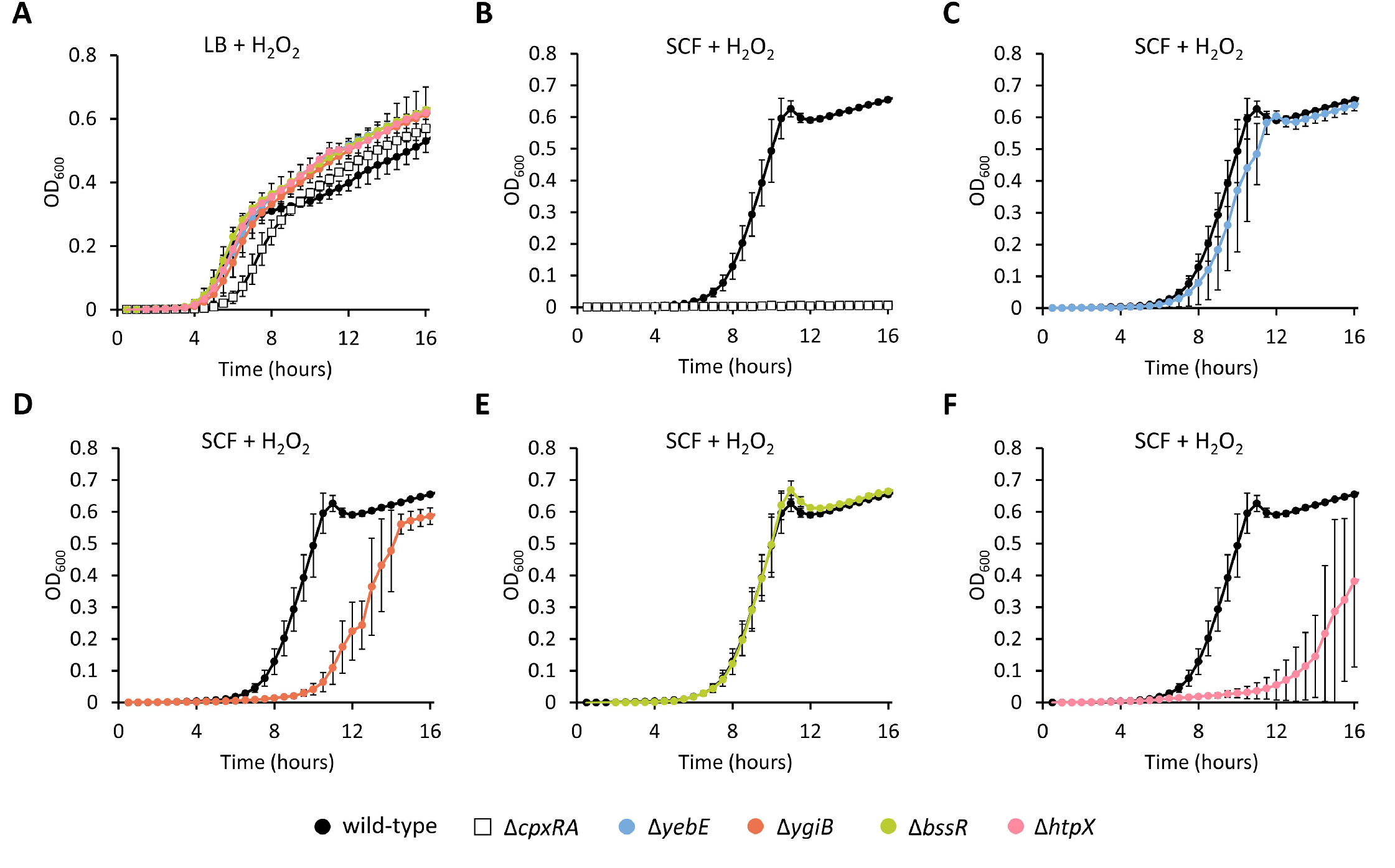
SCF with sub-inhibitory levels of oxidative stress inhibits Δ*cpxRA* mutant growth and highlights varying fitness defects in Δ*yebE*, Δ*ygiB*, and Δ*htpX* cells. Overnight cultures were standardized to an OD_600_ of 1.0 then subcultured 1:100 in (A) LB with 1mM H_2_O_2_ or (B-F) SCF with 1mM H_2_O_2,_ both buffered with 0.1M MOPS. Data points represent the mean of three biological replicate cultures with error bars indicating standard deviation.

Following these observations, we hypothesized that a contributing factor to the inability of the Δ*cpxRA* mutant to grow in SCF with an unstable pH could be attributed to the additive effects of reducing the expression of Cpx-regulated genes including *yebE, ygiB*, and *htpX*. To test this, double and triple mutants were generated using allelic exchange and grown in LB, SCF, and SCF with no MOPS (Figure 6). These mutants exhibited no aberrant growth phenotypes in LB (Figure 6A). In SCF, the double mutants containing Δ*ygiB* as well as the triple mutant had extended lag phases like that of Δ*cpxRA* (Figure 6B). These growth defects were exacerbated in unbuffered SCF where the only mutant that could successfully reach stationary phase was Δ*yebE*Δ*htpX*, albeit with an extended lag phase relative to wild-type cells (Figure 6C). The Δ*ygiB*Δ*htpX* double mutant as well as the triple mutant could not grow in unbuffered SCF (Figure 6C). These results support the hypothesis that the growth defects of the Δ*cpxRA* mutant in both buffered and unbuffered SCF can be attributed to the cumulative reduction in expression of Cpx-regulated genes, with *ygiB, yebE*, and *htpX* being especially important.

**Figure 6.**
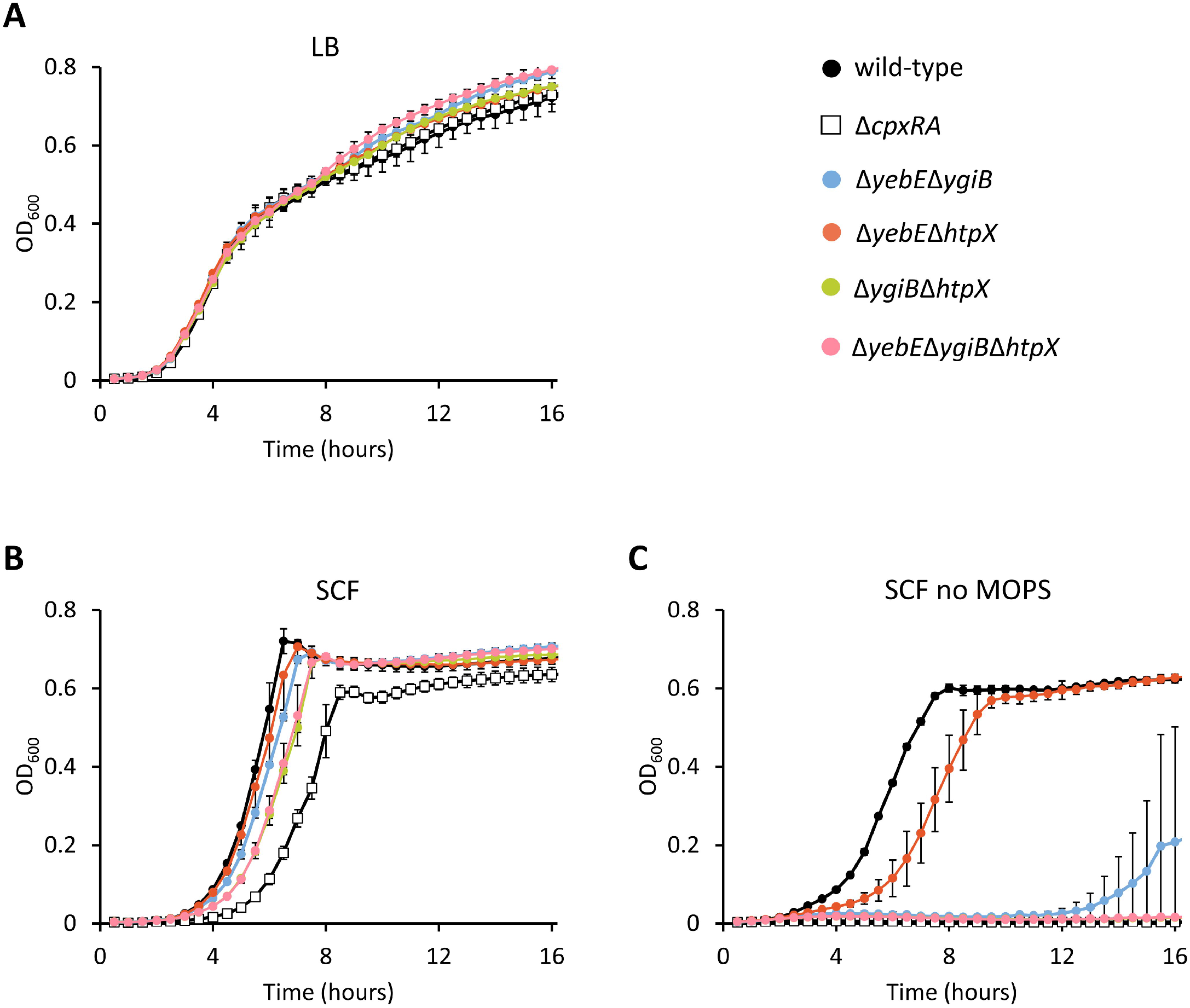
*C. rodentium* double and triple mutants of Cpx-regulated genes have similar fitness defects to Δ*cpxRA* cells. Overnight cultures were standardized to an OD_600_ of 1.0 then subcultured 1:100 into (A) LB, (B) SCF, and (C) SCF without MOPS buffer. Data points represent the mean of three biological replicate cultures with error bars indicating standard deviation.

To further analyze the fitness defects associated with the single-deletion mutants, *in vitro* competitive assays with a fluorescent wild-type *C. rodentium* strain, DBS100-AC, were conducted in LB and SCF. Of the five mutants tested, the Δ*cpxRA* and Δ*ygiB* mutants were significantly outcompeted by DBS100-AC after 24 hours growth in SCF (*P* < 0.05) (Figure 7). Importantly, this competitive disadvantage was not seen in LB, further supporting the use of SCF to investigate mutant phenotypes in *C. rodentium*. Unexpectedly, the Δ*cpxRA* mutant significantly outcompeted DBS100-AC in LB after 24 and 48 hours of growth (*P* < 0.01 and *P* < 0.05) (Figure 7B). This result was repeated in multiple experiments using different Δ*cpxRA* mutant glycerol stocks to confirm validity. The reason for the opposite results of competitions in LB and SCF requires further investigation. In addition, while statistically insignificant, the proportion of the Δ*htpX* mutant in SCF was lower than that of DBS100-AC and this trend was repeated in distinct experiments (Figure 7F). Therefore, the *in vitro* data collected thus far indicates that the Δ*cpxRA* mutant, along with mutants carrying deletions of the Cpx-regulated genes Δ*ygiB* and Δ*htpX*, have growth defects and a competitive disadvantage in SCF.

**Figure 7.**
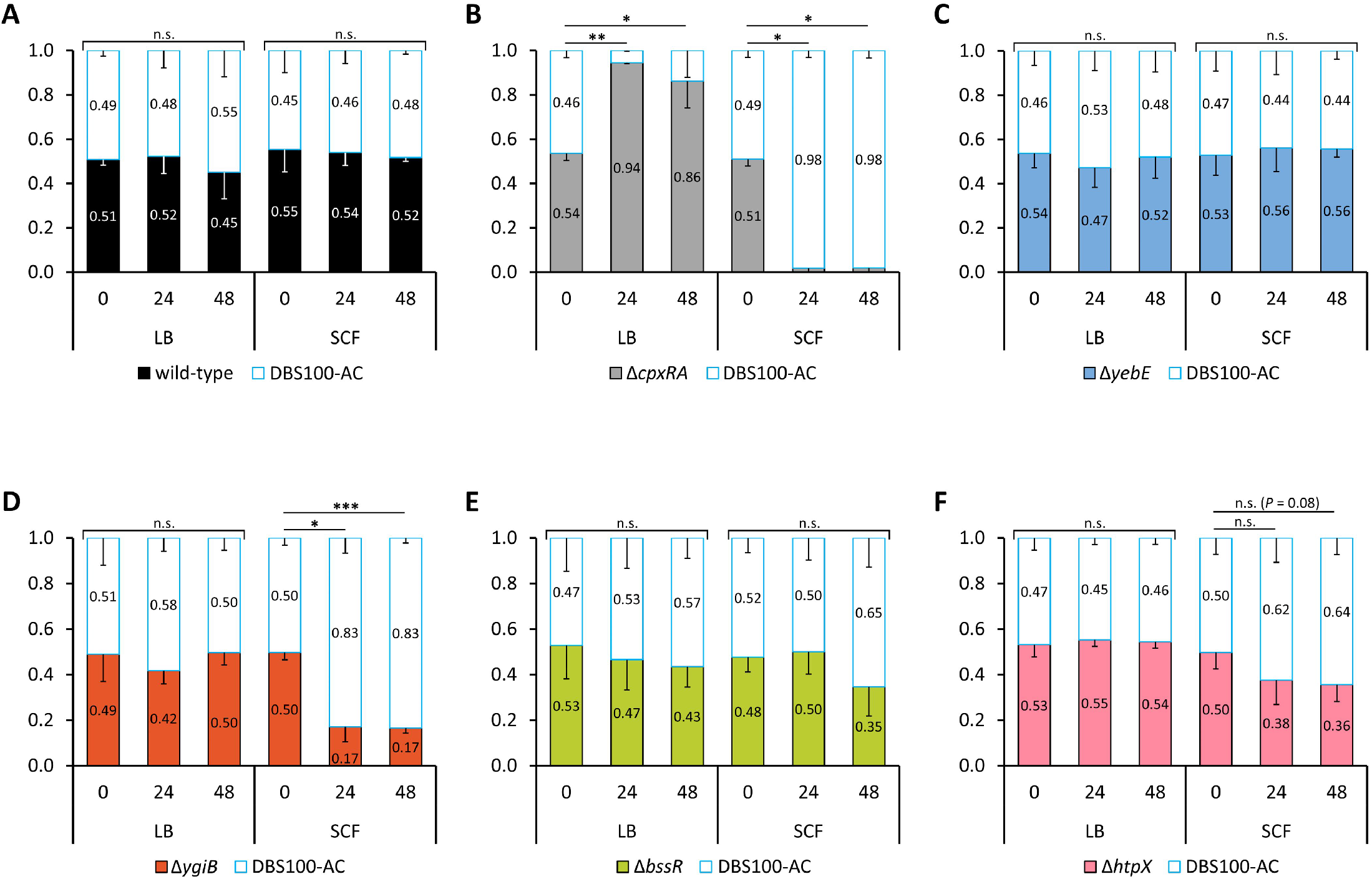
*In vitro* competitive assays in SCF indicate fitness defects in Δ*cpxRA*, Δ*ygiB*, and Δ*htpX* cells. Wild-type and mutant strains of *C. rodentium* were competed against a fluorescent *C. rodentium* strain expressing *amCyan* from a tetracycline inducible promoter. Strains were inoculated 1:1 into either LB or SCF and cultures were enumerated at 0, 24, and 48 hours on LB agar containing 0.2 ug/ml anyhrotetracycline. Bars represent the mean relative proportion of three biological replicate cultures with error bars indicating standard deviation. Significance was calculated using arcsine transformed data for 0 vs 24 hours and 0 vs 48 hours in both LB and SCF (\**P* < 0.05, \*\**P* < 0.01, \*\*\**P* < 0.001, n.s. not significant, Student’s *t*-test).

### In vivo *colonization levels replicate growth trends observed in SCF*

With *yebE, ygiB, bssR*, and *htpX* expression confirmed to be upregulated by the presence of CpxRA, we then tested whether these genes were required for the colonization or virulence of *C. rodentium* which could provide a possible explanation for why the removal of *cpxRA* is detrimental to pathogenesis (11–13). C57Bl/6J mice, which experience a self-limiting form of disease, were used to determine whether colonization was negatively impacted by any of the mutants (7). Only the Δ*cpxRA* mutant cells could not consistently colonize to the same level as wild-type cells and the other mutant strains (Figure 8A and Figure S6) (Day 4; *P* < 0.05, Day 9 and 12; *P* < 0.05, *P* < 0.01; Mann-Whitney U Test). While the Δ*yebE* mutant exhibited a slight lag in colonization levels on day 9 (*P* < 0.05) and the Δ*bssR* mutant showed a moderate increase in colonization of the colon on day 12 (*P* < 0.05), these minor statistical significances are not reflected in the degree of disease-state measured using a splenic index (Figure S6, Figure 8B). All strains caused a similar level of disease relative to wild-type except for Δ*cpxRA*, which had a significantly lower splenic index indicating attenuated virulence, thus confirming the results of previous studies (Figure 8B) (11, 12). Similarly, in C3H/HeJ mice, used for testing disease progression and survival, only the Δ*cpxRA* mutant exhibited a colonization defect as seen by an approximate 2-fold reduction in CFUs for three mice and undetectable levels in two on day 4 post-infection (Figure 8C). In addition, all five mice infected with the Δ*cpxRA* mutant survived until day 30 indicating a significant attenuation of virulence (Figure 8D). Importantly, the *in vivo* results shown here reflect similar trends to the growth phenotypes of the *C. rodentium* strains observed in SCF further indicating the importance of using physiologically relevant growth conditions in a laboratory setting prior to conducting animal trials.

**Figure 8.**
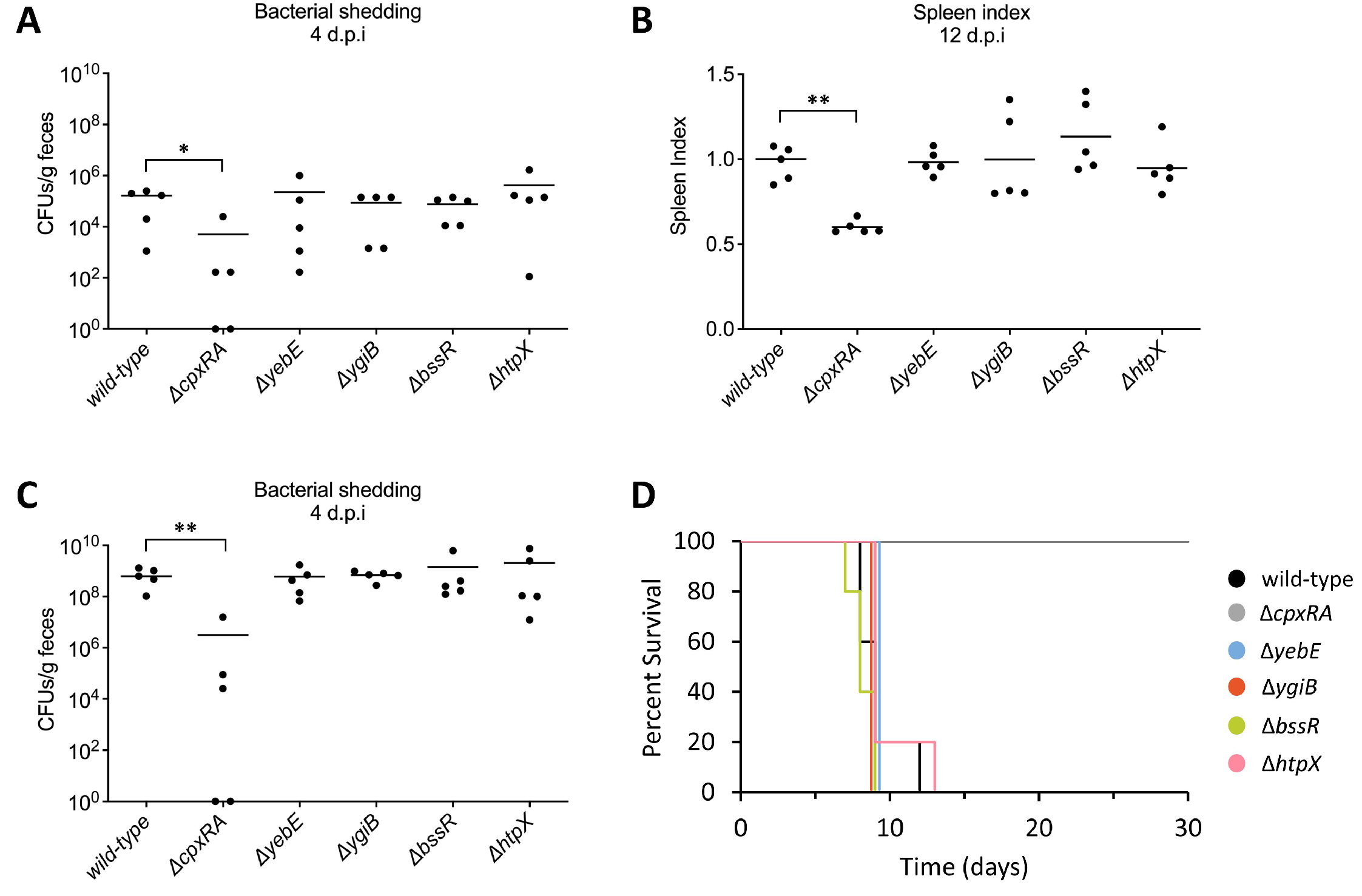
Genes of interest do not significantly contribute to *C. rodentium* colonization or virulence in C57Bl/6J or C3H/HeJ mice. (A) Colonization measured in colony forming units (CFUs) per gram of feces at 4 days post-infection and (B) spleen index in C57Bl/6J mice infected with *C. rodentium* mutants. (C) Colonization (CFUs/g) at 4 days post-infection in C3H/HeJ mice infected with *C. rodentium* mutants. (A-C) Horizontal lines indicate the median of n=5 mice and the asterisks indicate a statistically significant difference between mutant strain and wild-type cells (*\*P*<0.05, *\*\*P*<0.01, Mann-Whitney U Test). (D) Percent survival of C3H/HeJ mice infected with *C. rodentium* mutants. Mice were euthanized if any one of the following critical endpoints were reached: 20% body weight loss, hunching and shaking, inactivity, or body condition score of <2.

## Discussion

A frequently used gastrointestinal fluid is simulated gastric fluid (SGF) which has demonstrated the acid-tolerance of pathogens like EPEC, EHEC, *Vibrio cholerae, Listeria monocytogenes*, and *Salmonella* (42–45). Meanwhile, SCF is rarely, if ever, used in individual pathogen fitness studies as it has primarily been used for determining drug solubility and delivery systems which involve looking at the microbiota’s affect on drug release (46, 47). On the other hand, HG-DMEM is more commonly used in pathogen studies and has been used to induce virulence gene expression in *C. rodentium* prior to conducting animal model experiments (38, 39). In this study, we chose to investigate the role of the Cpx ESR and Cpx-regulated genes in a medium relevant to the conditions present in the colon by using SCF with the goal of identifying factors contributing to the Δ*cpxRA* mutants’ attenuation *in vivo* (11).

Throughout this study, we identified numerous growth conditions which influenced the activity of the Cpx ESR. The Cpx ESR is associated with monitoring proper membrane protein biogenesis, the repression of virulence factors, and maintaining cell wall integrity (16). A previous study published microarray data indicating that HG-DMEM moderately induced *cpxP* gene expression in EPEC overexpressing NlpE though they did not comment on this result being DMEM-dependent (41). Whilst DMEM has been used in previous Cpx-related studies to induce virulence gene expression, to our knowledge this is the first study to distinctly demonstrate a strong induction of the Cpx ESR in HG-DMEM and SCF relative to LB (13, 25, 26, 36, 37, 41). The activation of the Cpx ESR in HG-DMEM and SCF highlights the importance of stringent regulation of envelope functions under virulence factor expressing and physiologically relevant conditions.

Not only was the Cpx ESR activated in HG-DMEM and SCF, but it also had unique activity patterns demonstrated over time as well as was differentially impacted by shaking versus static conditions in a media-dependent manner. In *Salmonella enterica*, it has been shown that the expression of *hilA* is CpxA-dependent in low pH and it’s expression is increased when cultures are grown statically (48). In uropathogenic *E. coli*, the agglutination titer was opposite for wild-type and strains lacking the small RNA, RyfA, when grown shaking or statically in LB and human urine (49). Our observations of Cpx-regulated gene expression over time in different medias indicate the dynamic nature of the Cpx ESR and how the integration of numerous, likely intrinsic and extrinsic, signals can alter its activity. While single time points can indicate activation or repression by the Cpx ESR, collecting data over time allows for a further understanding of the reliance of gene expression on the Cpx ESR throughout the various phases of growth. Therefore, our study highlights the importance of culture conditions on the activity of the Cpx ESR and prompts questions as to the nature of the envelope stressors in the medias tested and by extension how those conditions could impact the physiology of growing cells.

### *The presence of the Cpx ESR negatively impacts the expression of master LEE regulator*, ler

Previous studies have shown that the Cpx ESR negatively impacts the expression of virulence factors. In EPEC, overexpression of the response regulator CpxR reduced the activation of LEE1, LEE4, and LEE5 while the removal of CpxR resulted in increased expression of all five LEE operons when in a non-pathogenic *E. coli* background that lacked *ler* indicating the regulation of LEE by the Cpx response was likely *ler-*independent (37). In EHEC, it has also been shown that the Cpx ESR negatively regulates virulence factors, including those that are LEE-encoded through Sigma factor 32 and the Lon protease (36). On the other hand, a recent study by Kumar et al. (50) proposed a model for EHEC suggesting that CpxR upregulates the expression of *ler* directly and that serotonin is an inhibitor of the Cpx ESR which results in reduced transcription of LEE. The authors used qRT-PCR and growth in LG-DMEM to show that *ler* was reduced in the absence of CpxR in EHEC and was similarly impacted in a *C. rodentium* Δ*cpxA* mutant. Due to these findings, we wished to verify the impact of the Cpx ESR on the expression of *ler* to determine whether the avirulence associated with the Δ*cpxRA* mutant could be due to reduced expression of the LEE in *C. rodentium*. Unlike Kumar et al. (50), our data indicates that in static SCF, as well as both high- or low-glucose DMEM, in static and shaking conditions, the expression of *ler* is consistently higher in the Δ*cpxRA* mutant (Figure 2, Figure S2). Therefore, our data suggests the colonization and virulence phenotypes observed for the Δ*cpxRA* mutant is not due to reduced expression of *ler* but perhaps could be from the overexpression of *ler* which may contribute to reduced fitness and inappropriately timed virulence mechanisms *in vivo*.

### *SCF is beneficial for determining susceptibilities of mutants to stressors and predicting colonization defects* in vivo

In this study we show that simulating the conditions *C. rodentium* cells face during colonization of the colon using SCF was able to uncover growth defects and susceptibilities that would have been unidentified in LB or HG-DMEM. Of note, wild-type *C. rodentium* grew to a significantly higher OD in SCF relative to HG-DMEM and had an increased growth rate relative to that in LB suggesting that this media simulates an environment this gastrointestinal pathogen has adapted to. Furthermore, mutants baring mutations that ablated part or all of the Cpx ESR grown in SCF were more susceptible to extraneous stressors than wild-type *C. rodentium*, and these phenotypes were exacerbated in double and triple mutants, as well as in competition assays. Thus, employing conditions that more closely emulate those of the intestinal lumen allowed us to uncover phenotypes for mutations in genes that likely play roles *in vivo* that might be missed in less sensitive infection models.

Interestingly, the Δ*cpxRA* mutant had an increased lag phase when grown in the presence of oxidative stress in LB while growth was abolished in SCF with H_2_O_2_ (Figure 5A and B). As detailed thoroughly in recent reviews, it is understood that *C. rodentium* utilizes aerobic respiration to outcompete host microbiota during colonization (6, 51, 52). A previous study found that deletion of the *cydAB* genes in *C. rodentium*, which are required for aerobic respiration, resulted in a severe reduction of growth *in vivo* (53). Following this, it was determined that disruption to the mitochondrial respiration of intestinal epithelial cells is largely responsible for *C. rodentium* infection causing oxygenation of the mucosal surface as opposed to solely colonic crypt hyperplasia (54, 55). The Cpx ESR has been implicated in the regulation of aerobic respiration in EPEC where it has been shown that removal of *cpxRA* reduced the oxygen consumption capabilities of cells which was attributed to problems with cytochrome *bo*_*3*_ oxidase biogenesis or function (56). Similar conclusions have been made in *Salmonella* Typhimurium which also utilizes aerobic respiration to expand in the gastrointestinal tract and experiences colonization defects in the absence of CpxRA (57, 58).

A follow up study to Lopez et al. (53) found that prior to expansion by aerobic respiration, *C. rodentium* utilizes host derived H_2_O_2_ as an electron acceptor during anaerobic respiration via cytochrome *c* peroxidase (*ccp*) (59). Both wild-type and Δ*ccp* mutant cells could survive in LB in the presence of mM concentrations of H_2_O_2_, though in the absence of *ccp* there was increased expression of the catalase-peroxidase, *katG*, suggesting that cells were experiencing higher levels of oxidative stress. The RNA-seq data in this study also showed *katG* expression was significantly induced in the Δ*cpxRA* mutant grown in HG-DMEM which suggests the cells were experiencing elevated levels of oxidative stress (13). Given that rapid expansion by aerobic respiration is a proposed mechanism of *C. rodentium* pathogenesis in overcoming colonization resistance and the production of reactive oxygen species by intestinal epithelial cells is an important defense mechanism, our results support the hypothesis that the Cpx ESR is required to successfully adapt to oxidative stressors during colonization in the gastrointestinal tract (53, 60–62).

Previous research using variations of simulated gut media have indicated specific bacterial responses to conditions emulating the gastrointestinal tract. Musken et al. (63) used simulated intestinal fluids mimicking the ileum and colon to demonstrate differential expression of a major fimbrial subunit in sorbitol-fermenting EHEC while Polzin et al. (64) found that EHEC proteins involved in nucleotide biosynthesis and the expression of Shiga toxins were increased in simulated ileal and colonic environments. In addition, it was found that outer membrane vesicle (OMV) production and cytotoxicity, which is an important virulence factor as well as OMV-associated Shiga toxin 2a in EHEC, was increased in both simulated ileal and colonic environments (65). On the other hand, OMV cytotoxicity and OMV-associated Shiga toxin 2a was not increased in DMEM which was confirmed with RT-qPCR for *stx*_*2a*_ expression (65). One study in *Salmonella* demonstrated differences in cell viability upon exposure to gastrointestinal fluids where they found gastric juice with a pH of 4 or 5 in conjunction with bile salts from simulated intestinal juice reduced cell viability greater than acidic pH alone (45). These studies, in conjunction with the data presented here, highlight the importance of using a physiologically relevant environment when it comes to monitoring gene and protein expression in cells during gastrointestinal survival, colonization, and virulence.

## Acknowledgements

The authors thank the University of Alberta Molecular Biology facility for assisting with Sanger sequencing and Kat Pick for finding and suggesting the recipe of simulated colonic fluid used as well as for thoughtful discussion of results.

This research was supported by operating grants by the National Sciences and Engineering Research Council (NSERC) to TR and to SG.

